# Spatial geometry-aware deep learning for deciphering tissue structure from spatially resolved transcriptomics

**DOI:** 10.1101/2025.05.11.652625

**Authors:** Xingyi Li, Xiangting Jia, Dongmin Zhao, Jialuo Xu, Gaoyuan Du, Yang Qi, Yiqi Chen, Yingfu Wu, Junnan Zhu, Feng Wei, Xuequn Shang

**Affiliations:** School of Computer Science, Northwestern Polytechnical University, Shannxi, China; State Key Laboratory of Multimodal Artificial Intelligence Systems, Institute of Automation, CAS, Beijing, China; Department of Hepatobiliary and Pancreatic Surgery, General Surgery Center, First Hospital of Jilin University, Jilin, China

**Keywords:** Spatial transcriptomics, spatial geometry-aware, Autoencoder, Geometric graph learning

## Abstract

Recent advances in spatially resolved transcriptomics have enabled high-throughput gene expression profiling while preserving spatial context, thereby facilitating the investigation of spatial heterogeneity within tissues. Here, we present SpatialGEO, a spatial geometry-aware deep learning framework designed to decipher tissue organizational structures through dual-encoder feature extraction and geometric graph learning. Experimental results demonstrate the superior accuracy of SpatialGEO in identifying spatial regions across datasets encompassing diverse biological contexts and spatial resolutions. Moreover, SpatialGEO uncovers key mechanisms of immune evasion and regulation within the tumor microenvironment and provides novel biological insights into mouse embryonic development.

## 1 Introduction

Deciphering tissue structure and cellular heterogeneity is essential for understanding how complex biological functions emerge from coordinated interactions among diverse cell types within organized spatial contexts. Spatial transcriptomics techniques not only capture gene expression but also preserve the spatial locations of cells within tissues, providing an unprecedented opportunity to study the spatial correlations between tissue structure and function.

In recent years, spatial transcriptomic technologies have rapidly evolved, primarily encompassing in situ capture-based methods (e.g., 10x Visium [1], Slide-seq [2, 3], and Stereo-seq [4]) and fluorescence imaging-based methods (e.g., MERFISH [5, 6], osmFISH [7], and seqFISH+ [8]). These techniques continue to advance in spatial resolution and gene coverage, providing powerful tools for deciphering tissue architecture and spatial heterogeneity.

The emergence of spatial transcriptomics has enabled unprecedented exploration of spatial heterogeneity in tissue microenvironments. However, a key challenge remains: how to effectively integrate gene expression with spatial information. Early methods such as K-means, Louvain, and Seurat relied solely on gene expression similarity, neglecting spatial context and yielding fragmented, biologically implausible domains. Statistical models like Giotto [9] and BayesSpace [10] improved spatial coherence by incorporating spatial priors and neighborhood correlations, but are limited by rigid neighborhood assumptions and high computational costs. Machine learning methods including stLearn [11], UTAG [12], and SpaGene [13] integrate spatial features via feature engineering, yet depend on shallow, handcrafted representations that fail to capture complex nonlinear spatial patterns. Graph neural network (GNN)-based models, such as SpaGCN [14], model spatial relationships through graph structures but are often constrained by limited message-passing capacity. More recent self-supervised approaches like GraphST [15] and ConST [16] enhance representation learning via contrastive strategies, though their performance may suffer from noise or biases in neighborhood definitions, leading to semantic ambiguity. Recent deep learning approaches based on autoencoder, such as STAGATE [17] and SEDR [18], have demonstrated strong potential in embedding spatial transcriptomic data and identifying spatial domains. However, most of these methods rely on a single Graph Autoencoder (GAE) architecture, which primarily captures local topological structures based on predefined spatial graphs, dependent on the quality of the input graph. The graph structure is often a simple, discrete topology that does not accurately reflect the complex spatial relationships within the tissue organization.

To overcome these limitations, we propose the SpatialGEO model, which adopts a heterogeneous dual-source encoder architecture to extract features and introduces a novel geometric-aware cell embedder to incorporate distance matrices for modeling continuous geometric relationships among cells or spots, supplementing the lack of spatial continuity in standard graph data and overcoming the limitations of discrete graph topology. This significantly improves the model’s ability to perceive spatial information in tissue structures and provides strong support for more comprehensive and accurate spatial modeling. We extensively evaluated the SpatialGEO model across various spatial transcriptomics platforms with different spatial resolutions, such as 10x Visium, Stereo-seq, Slide-seqV2, MERFISH, and osmFISH. Experimental results demonstrate that SpatialGEO accurately identifies spatial domains and reveals spatial heterogeneity across diverse datasets. The model shows exceptional generalization and robustness, providing valuable insights into tissue development and disease mechanisms.

## 2 Results

### 2.1 Overview Of SpatialGEO

The raw gene expression matrix is standardized, and a spatial graph is constructed using the k-nearest neighbors (KNN) algorithm to capture local spatial relationships among spots. SpatialGEO treats gene expression profiles and the spatial graph as two heterogeneous data sources. To model these data jointly, it employs a dual-encoder architecture comprising an autoencoder (AE) and a graph autoencoder. The AE captures intrinsic gene expression patterns, while the GAE learns the tissue’s spatial organization. The outputs of both encoders are fused into a unified representation that integrates gene expression and spatial information. To enhance spatial modeling, SpatialGEO introduces a geometric-aware cell embedder module. This module performs multi-hop message passing to aggregate contextual information from neighboring spots, thereby improving spatial awareness. Additionally, a geometric graph learning mechanism is applied to model continuous spatial relationships, overcoming the limitations of discrete graph structures. These enhancements are combined to produce a final latent representation within a unified embedding space. And an adaptive target distribution strategy is integrated into the training process to align gene expression and spatial features effectively. The resulting integrated representation supports diverse downstream analyses, including unsupervised clustering to delineate spatial domains and decoding to reconstruct denoised gene expression profiles for regulatory and functional analysis.

### 2.2 Quantitative evaluation of SpatialGEO on brain datasets

We applied SpatialGEO to human and mouse brain datasets to quantitatively evaluate the effectiveness of the method. The human dorsolateral prefrontal cortex (DLPFC) dataset, generated using the 10x Genomics Visium platform, contains 12 slices from three independent neurotypical adult donors [19]. Each section includes six cortical layers (L1–L6) and white matter (WM) with well-defined morphological boundaries. Maynard et al.[19] manually annotated the cortical layers and WM of each slice according to morphological characteristics and genetic markers, and these annotations were used as ground truth for the evaluation of spatial clustering methods (Fig. 2A). To systematically evaluate and compare the performance of different clustering algorithms for spatial domain identification in the DLPFC, we used k-means clustering on gene expression data as a baseline and selected six state-of-the-art algorithms for comparison: SpaGCN, stLearn, DeepST, SEDR, STAGATE, and GraphST. We performed clustering analysis on slice 151672 (Figure 2A) using all eight methods.

**Fig. 1.**
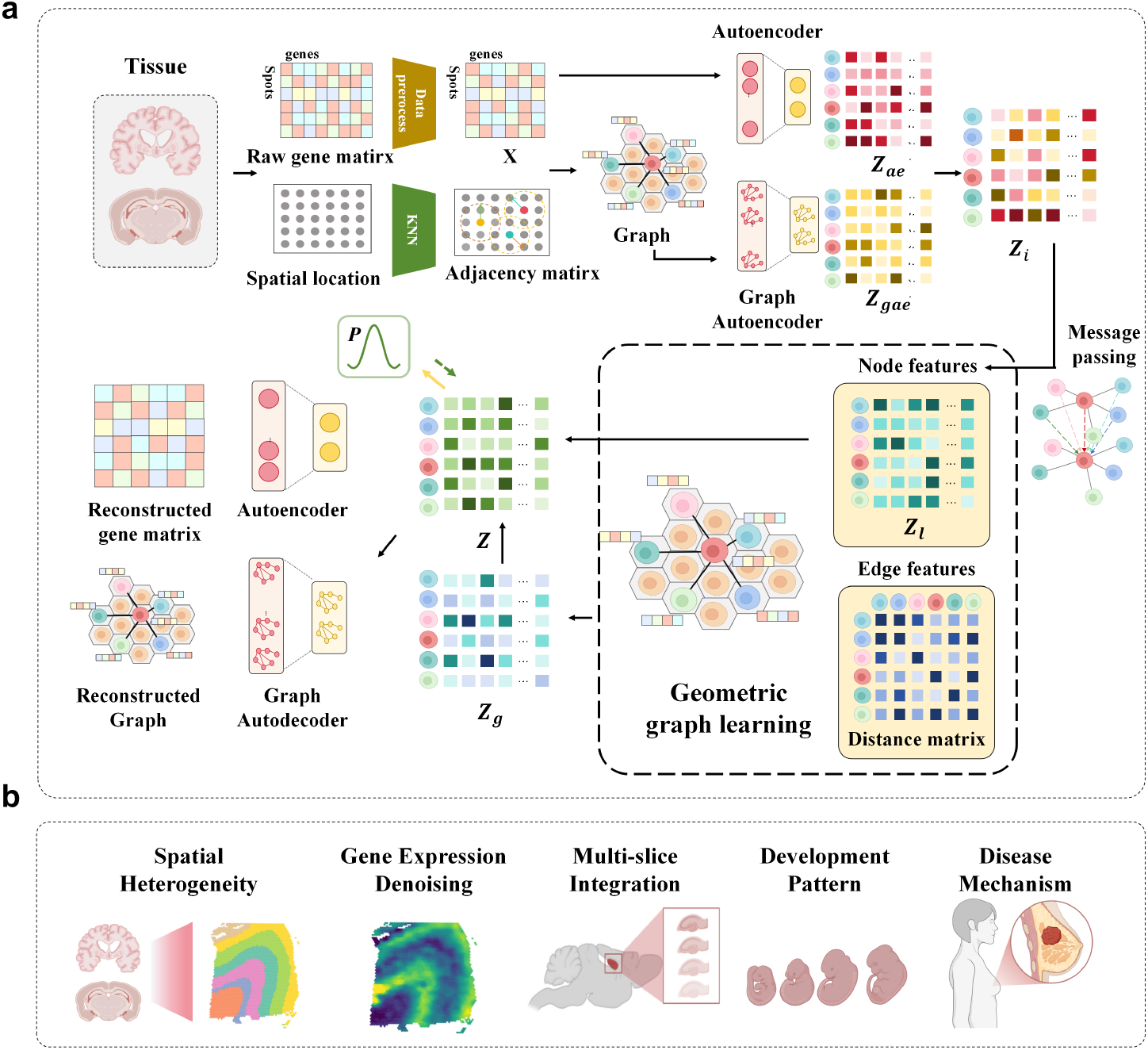
Overview of SpatialGEO framework. (**A**) **Heterogeneous Dual-Source Encoders architecture**: Raw gene expression matrix (spots × genes) is preprocessed and combined with spatial coordinates to construct a KNN graph. Heterogeneous dual-source encoders (Autoencoder for gene patterns and Symmetric Graph Autoencoder for spatial topology) generate embeddings *Z*_*ae*_ and *Z*_*gae*_, dynamically fused into *Z*_*i*_. (**B**) **Geometric-aware cell embeddor**: Multi-hop message passing on *Z*_*i*_ produces *Z*_*l*_, followed by geometric graph learning (distance matrix replaces KNN edges) to refine *Z*_*g*_. Skip-connection integrates *Z*_*l*_ (local) and *Z*_*g*_ (global) into final embedding *Z*. (**C**) **Downstream tasks**: *Z* enables spatial domain identification, trajectory analysis, batch integration, gene denoising, and regulatory network inference.

**Fig. 2.**
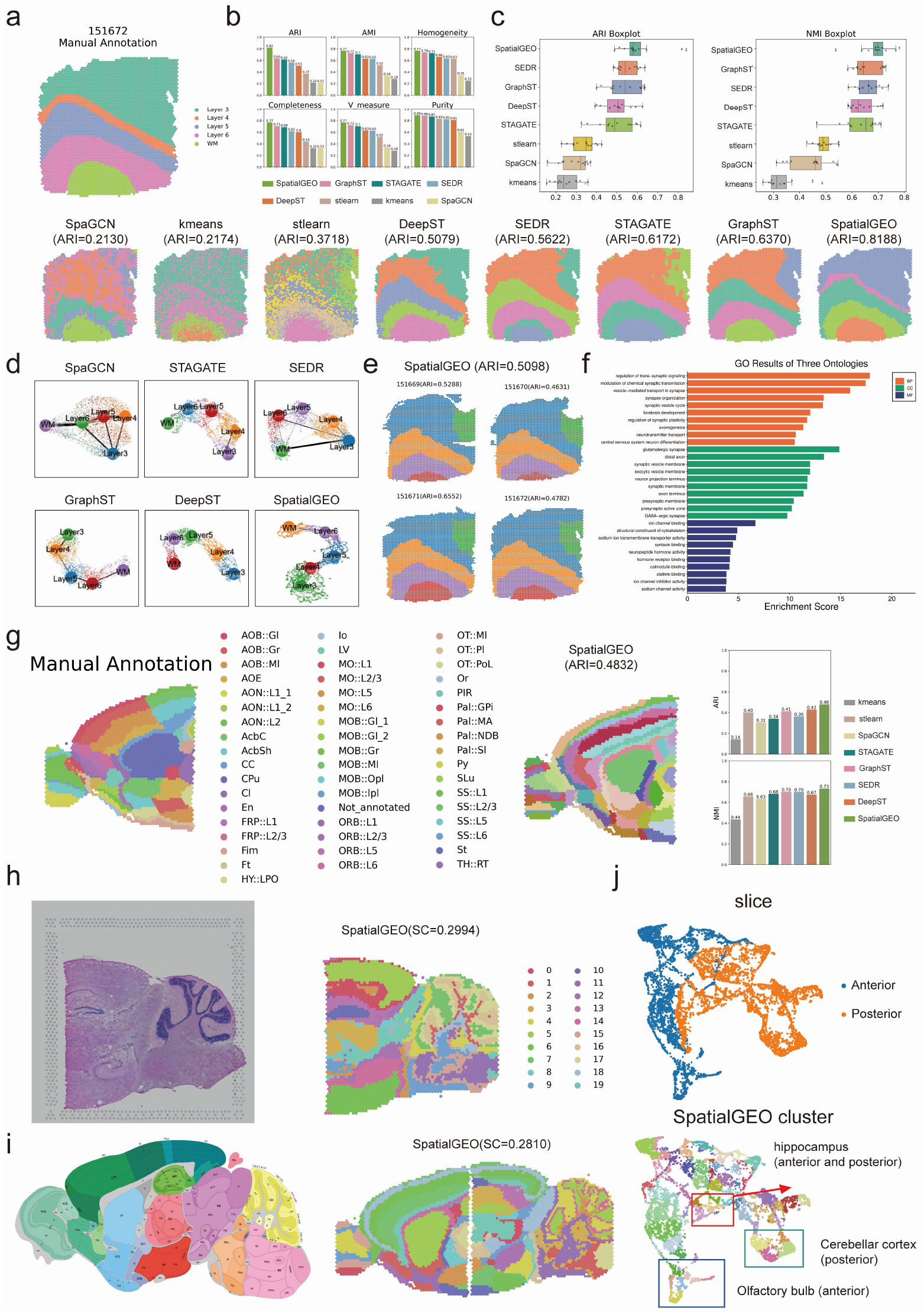
Comparison of clustering methods on Brain datasets. **(A)** Manual annotation and clustering results of sample #151672 from the human DLPFC dataset. Eight methods are compared: SpaGCN, k-means, stLearn, DeepST, SEDR, STAGATE, GraphST, and SpatialGEO.**(B)** Clustering performance evaluation on sample #151672 using six metrics: Adjusted Rand Index (ARI), Adjusted Mutual Information (AMI), Homogeneity, Completeness, V-measure, and Purity.**(C)** Boxplots showing ARI (left) and NMI (right) across all 12 slices of the DLPFC dataset for the eight clustering methods.**(D)** PAGA trajectory analysis results on sample #151672 for all methods.**(E)** Visualization of spatial clustering results (SpatialGEO) after horizontal integration of four slices: #151669, #151670, #151671, and #151672.**(F)** GO enrichment analysis of Layer 4 and Layer 5 identified by SpatialGEO in sample #151672.**(G)** Manual annotation and SpatialGEO clustering results on the mouse brain forebrain region. Right: ARI and NMI performance comparison of eight methods.**(H)** Histological image of the mouse brain hindbrain and SpatialGEO clustering visualization.**(I)** Allen Brain Atlas annotation (left) and SpatialGEO clustering results (right) across the whole mouse brain.**(J)** UMAP visualization of two mouse brain slices, showing spatial clusters identified by SpatialGEO in anterior/posterior regions including hippocampus, cerebellar cortex, and olfactory bulb.

Clustering by K-means based solely on gene expression resulted in unclear anatomical structures. Similarly, the results from SpaGCN and stLearn appeared mixed and could not effectively identify different anatomical structures. In contrast, the four deep learning algorithms (DeepST, SEDR, STAGATE, GraphST) improved clustering performance but still exhibited issues such as intra-layer fragmentation, especially within Layer 3, and unclear layer boundaries. Remarkably, SpatialGEO achieved outstanding performance, clearly delineating layer boundaries without producing outlier clusters. SpatialGEO’s segmentation was highly consistent with the manual annotations, particularly successfully identifying the Layer 4 region. We quantitatively evaluated the model performance through the adjusted Rand Index (ARI)[20]. SpatialGEO achieved 0.8188 in this metric, significantly outperforming the second-ranked GraphST method (ARI = 0.6370), with an improvement of nearly 28% (Fig. 2B). To comprehensively measure the consistency between the prediction results and the true labels, we further introduced multiple indicators such as adjusted mutual information (AMI), Homogeneity, Completeness, V-Measure and Purity for evaluation. The results show that SpatialGEO achieved the highest score in all metrics, indicating its outstanding clustering performance.

We further assessed the robustness of the model across all 12 DLPFC slices (Fig. 2C). SpatialGEO demonstrated consistently strong performance, achieving the highest median ARI (0.580) and the smallest interquartile range (IQR = 0.053), highlighting its superior generalization capability. Additionally, it attained the highest median NMI score (0.713) [21], further confirming its robustness and consistency across diverse samples. Furthermore, using UMAP dimension reduction combined with PAGA topology analysis [22, 23], SpatialGEO successfully reconstructed a trajectory from WM through Layer 6 to Layer 3 (Figure 2D). Other methods like SpaGCN, SEDR, GraphST, and DeepST produced redundant branches inconsistent with known anatomical paths. Although STAGATE correctly recognized topology, its distribution across layers was inaccurate. SpatialGEO was also applied to integrate four consecutive slices from one donor (#151669–#151672). After vertical integration and batch effect removal, SpatialGEO maintained clear clustering (average ARI = 0.51). (Figure 2E). To explore biological functions, we conducted Gene Ontology (GO) enrichment analysis on Layer 4 and Layer 5 differentially expressed genes identified by SpatialGEO (Figure 2F). Layer 4 genes were enriched in synaptic signaling regulation and plasticity, suggesting a role in working memory storage. Layer 5 genes were enriched in axonogenesis and synaptic vesicle cycle processes, consistent with long-range projection and behavior output functions.

In the mouse anterior brain tissue dataset (Fig. 2G), SpatialGEO accurately delineated anatomically meaningful regions, including the olfactory bulb and dorsal pallium, achieving the highest ARI (0.48) and NMI (0.73) when using manual annotations from Long et al.[15] as the reference. In particular, within the olfactory bulb (OB) region, SpatialGEO successfully reconstructed the laminar structure of the main olfactory bulb (MOB), exhibiting high concordance with the Allen Mouse Brain Atlas [24]. Furthermore, key subregions of the striatum—caudoputamen, nucleus accumbens, and olfactory tubercle—were distinctly identified, which none of the other comparative methods could achieve. For the unlabeled posterior brain dataset (Fig. 2H), we evaluated performance using the Silhouette Coefficient (SC) [25] and Davies–Bouldin Index (DB) [26]. Although K-means showed a DB value closer to 1, its clustering results were highly fragmented and failed to capture coherent spatial domains. In contrast, SpatialGEO achieved a better clustering quality with a DB score of 1.23, as evidenced by the clear delineation of the hippocampus and cerebellum regions. Notably, the spatial organization of the cerebellar region closely matched that of the Allen Brain Atlas. We further conducted a horizontal integration analysis across mouse brain slices (Fig. 2I). After aligning tissue sections and applying clustering (with the number of clusters fixed at 52 for consistency), we compared the results against anatomical annotations from the mouse brain atlas. SpatialGEO consistently produced clusters that closely matched known anatomical structures. Its spatial clusters showed strong colocalization with well-characterized brain architectures, including the dentate gyrus, CA3–CA1 axis, olfactory bulb, cerebellum, and hippocampus. Visualization using UMAP following integration (Fig. 2J) revealed that spatial embeddings of shared regions (such as cortex and hippocampus) from different slices exhibited partial overlap—rather than complete mixing—demonstrating SpatialGEO’s ability to mitigate batch effects while preserving biological structure. Slice-specific areas remained distinct, reflecting expected differences in functional regions across sections. These results underscore SpatialGEO’s robust capacity for spatial domain alignment and batch correction.

### 2.3 SpatialGEO enables efficient denoising of spatial transcriptome data

We performed clustering on the DLPFC slice #151507 and found that SpatialGEO achieved the highest clustering accuracy among all methods evaluated (ARI = 0.61; Fig. 3A). Denoised gene expression profiles were recovered using the decoder module of SpatialGEO. To assess its denoising performance, we applied SpatialGEO to the same slice and systematically compared the spatial expression patterns of canonical layer-specific marker genes before and after denoising. Notably, laminar enrichment of marker genes was markedly improved in the denoised data (Fig. 3B) [27, 28]. *HPCAL1*, a known marker of layer 2 (L2) excitatory neurons, exhibited strong and selective expression in L2 after denoising (median = 0.97), with minimal expression in other layers—effectively eliminating cross-layer signal contamination observed in the raw data. This pattern is consistent with its well-established layer specificity. Similarly, *PCP4*, a marker of layer 5 (L5) pyramidal neurons, showed a sharp expression peak in L5 (median = 0.51) that corresponded precisely to its anatomical localization. In the raw data, *ATP2B4* exhibited diffuse and ambiguous expression; however, denoising revealed a clear bimodal distribution, with distinct enrichment in both L2 (0.20) and L6 (0.09), thereby resolving its original spatial ambiguity. These results indicate that SpatialGEO effectively distinguishes true biological signals from noise-induced artifacts. The myelin-associated gene *MOBP* demonstrated robust aggregation in the white matter after denoising (median = 2.36), with suppressed expression across cortical layers (L1: − 0.31, L2: − 0.27, L3: − 0.28, L4: − 0.22, L5: 0.29, L6: 0.81). This spatial pattern aligns with the distribution of glial cells, underscoring the algorithm’s capacity to preserve biologically meaningful localization. *SNAP25*, a neuronal gene with broad laminar expression in the cortex, retained its expected pan-laminar profile post-denoising, while its expression in the white matter was substantially reduced (median = −0.75), further highlighting SpatialGEO’s ability to maintain biologically relevant gradients while removing nonspecific signal. Consistent denoising performance was also observed in 11 additional DLPFC slices, where SpatialGEO consistently improved the spatial specificity of gene expression patterns, as confirmed by gene expression visualization and stacked violin plots.

**Fig. 3.**
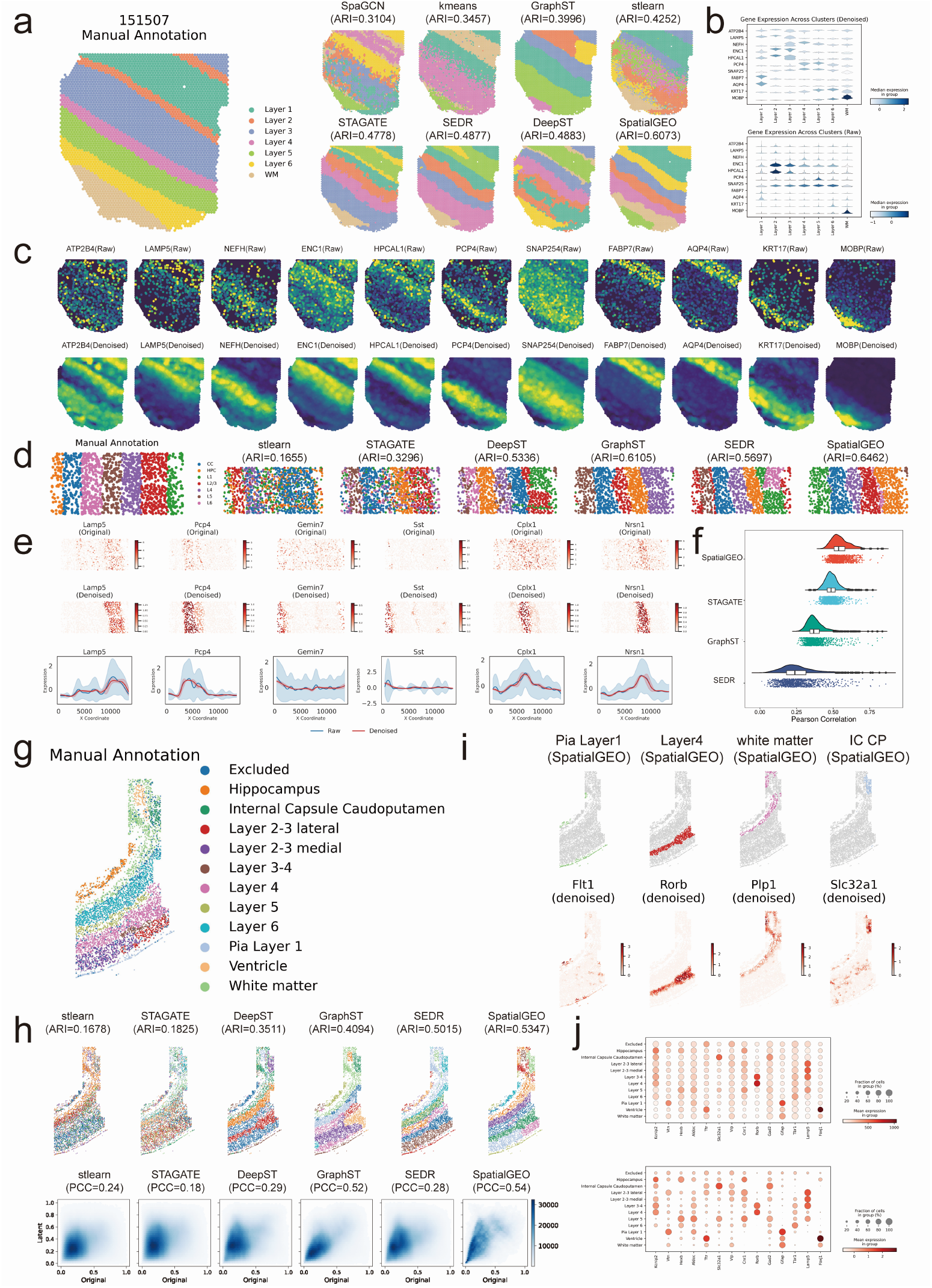
Benchmarking SpatialGEO across human and mouse brain datasets. (**a**) Manual annotation of DLPFC slice 151507 (left) and clustering results from eight methods, with SpatialGEO yielding the highest accuracy (ARI = 0.61). (**b**), Violin plots showing expression of layer-specific marker genes across clusters before (top) and after (bottom) denoising with SpatialGEO. (**c**), Spatial expression patterns of representative marker genes before (top row) and after (bottom row) denoising, highlighting enhanced laminar specificity post-denoising. (**d**), Reference annotation and clustering results of six methods on the mouse visual cortex dataset from the STARMAP platform. ARI values are indicated. (**e**), Spatial expression profiles of canonical layer-specific genes: raw (top), denoised by SpatialGEO (middle), and smoothed expression along the X-axis (bottom) showing denoising-enhanced laminar contrast. (**f**), Violin plots comparing recovery performance of 2,000 differentially expressed genes across four methods in the mouse cortex dataset, evaluated by Pearson correlation. (**g**), Manual annotation of the mouse somatosensory cortex dataset from the osmFISH platform. (**h**), SpatialGEO-recovered spatial domains and marker gene expressions (Fit1, Rorb, Plp1, Slc32a1) in selected regions (Pia layer 1, layer 4, white matter, internal capsule). (**i**), Clustering results and pair-wise distance correlation density plots (PCC, Pearson correlation coefficient) between original and imputed spaces across six methods. (**j**), Dot plots of layer-specific genes in raw (top) and denoised (bottom) data, where dot size indicates expression level and color intensity denotes expression strength. SpatialGEO markedly improves laminar resolution and expression specificity.

To evaluate the denoising capability of SpatialGEO, we applied it to STARmap transcriptomic data derived from the mouse visual cortex [29]. We first conducted unsupervised clustering on the spatial transcriptomics dataset. Among all evaluated methods, SpatialGEO achieved the highest Adjusted Rand Index (ARI) (Fig. 3d), demonstrating superior clustering performance and better preservation of biologically meaningful structures. We then investigated the spatial expression patterns of six specifically differentially expressed genes—*Lamp5, Pcp4, Gemin7, Sst, Cplx1*, and *Nrsn1* (Fig. 3e) [29, 30]. In the raw data, these genes exhibited noisy and indistinct spatial expression, limiting their ability to delineate cortical layers. After applying SpatialGEO, their expression patterns became markedly more structured and layered, consistent with known laminar architecture. To further quantify this improvement, we analyzed the expression gradients of these layer-enriched genes along the X-axis. Compared to the original data, the SpatialGEO-denoised expression profiles displayed smoother spatial variation (red line), clearer peak structures, and reduced variability, as evidenced by the narrower confidence bands (shaded red) versus the original (shaded blue). This suggests that SpatialGEO not only effectively attenuates technical noise but also preserves and enhances authentic biological signal, thereby improving interpretability and facilitating the identification of spatial gene expression domains. Finally, we compared the performance of SpatialGEO with other methods in preserving native gene expression trends using raincloud plots (Fig. 3f). SpatialGEO consistently achieved higher Pearson correlation coefficients between the denoised and raw expression data across most genes. Furthermore, the distribution of correlation values showed lower skewness and higher kurtosis, indicating tighter agreement with the original data and improved consistency across genes. Collectively, these results underscore the robustness and fidelity of SpatialGEO in denoising spatial transcriptomics data while maintaining biologically meaningful expression patterns, thereby facilitating downstream analysis of tissue architecture and cell identity.

We also collected a spatial transcriptomics dataset of the mouse somatosensory cortex acquired using the in situ hybridization fluorescence imaging technique osm-FISH [29]. The dataset comprises gene expression profiles of 4,839 cells across 33 genes. Ground truth annotations divide these cells into 11 cortical subregions, including Pia Layer 1, Layer 4, and white matter (Fig. 3g).To quantitatively assess clustering performance, we benchmarked six representative spatial transcriptomics analysis methods—stLearn, STAGATE, DeepST, GraphST, SEDR, and SpatialGEO—using the Adjusted Rand Index (ARI) as the evaluation metric. As shown in Fig. 3h, **SpatialGEO** achieved the highest ARI of 0.53, indicating substantially improved agreement with manual annotations over competing methods. Notably, **SpatialGEO** was the only method capable of accurately resolving the Layer 2/3 lateral subregion, underscoring its sensitivity to subtle spatial heterogeneity. Subsequently, we calculated the Euclidean distances between any two points in the latent spaces generated by each method, and conducted a correlation analysis between these distances and their corresponding distances in the original space, as shown in Figure 6b. The results indicated that the correlation between the latent space generated by SpatialGEO and the original space was significantly higher than that of other methods. The Pearson correlation coefficient (PCC) reached 0.54, demonstrating that it performed better in preserving the topological structure of the original space.

Denoising further enhanced the anatomical concordance of spatial expression patterns for canonical marker genes (Fig. 3i) [29]. For example, the endothelial cell marker *Flt1* was accurately restricted to Pia Layer 1, while the inhibitory neuronal marker *Slc32a1* was specifically enriched in the Internal Capsule Caudoputamen region, demonstrating robust cell type–specific spatial localization. The Layer 4 pyramidal neuron marker *Rorb* exhibited strong expression in the Layer 4 domain identified by SpatialGEO, highlighting its ability to resolve laminar cortical structures. Additionally, the oligodendrocyte-specific gene *Plp1* showed markedly higher expression in white matter than in other regions, indicating precise delineation of glial distributions. To further validate the effectiveness of denoising, we selected representative high-expression genes within each anatomical region—including key developmental markers (Fig. 3j) such as *Kcnip2, Vtn, Hexb*, and *Aldoc*—and visualized their raw and denoised spatial expression using dot plots. The denoised profiles revealed clearer stratified expression layers and stronger regional specificity, demonstrating the improved spatial resolution and enhanced interpretability of downstream analyses.

### 2.4 Unraveling the Tumor Microenvironment in Breast Cancer: From Healthy Tissue to Invasive Carcinoma

Tumor heterogeneity is a hallmark of cancer that complicates diagnosis, prognosis, and treatment, particularly in breast cancer (BRCA), where both spatial and molecular diversity within the tumor microenvironment profoundly influence disease progression and therapeutic response [31]. To unravel this complexity, we applied multiple clustering strategies to spatial transcriptomic data derived from human BRCA samples. Among these, the SpatialGEO method demonstrated superior performance, achieving the highest Adjusted Rand Index (ARI = 0.70; Figure 5a), and offered a precise delineation of tissue boundaries with strong concordance to manual annotations. Based on this result, we performed a focused analysis of four spatially distinct regions identified by SpatialGEO (Figure 5b): Cluster17 (healthy tissue), Cluster8 (ductal carcinoma in situ/lobular carcinoma in situ, DCIS/LCIS), Cluster14 (tumor margin), and Cluster2 (invasive ductal carcinoma, IDC). These domains were further investigated using the mfuzz clustering method, which enabled the identification of co-expressed gene clusters across spatial contexts. Subsequent functional enrichment analyses revealed distinct biological programs underpinning each region. Cluster17, corresponding to healthy tissue, was enriched for gene clusters involved in chemokine signaling and immune homeostasis (e.g., leukocyte chemotaxis, IL-1 response), as well as extracellular matrix (ECM) remodeling, suggesting a protective microenvironment characterized by both immune surveillance and structural integrity. In contrast, Cluster8, representing DCIS/LCIS, showed elevated expression of gene clusters associated with interferon-mediated antiviral responses and lipid metabolic reprogramming, including enhanced fatty acid oxidation, while concurrent MHC-II activation and potential regulatory T cell recruitment implied early-stage immune suppression despite preserved epithelial polarity. Cluster14, located at the tumor margin, exhibited molecular signatures indicative of both immune activation and evasion, with complement system engagement and humoral immune response potentially counteracted by upregulation of complement inhibitors such as CD55 and CD59. In parallel, increased ECM remodeling, including collagen crosslinking and FAK/PI3K signaling, suggested active epithelial-mesenchymal transition (EMT) processes facilitating tumor cell dissemination. Cluster2, representing IDC, was characterized by high expression of genes involved in cell cycle dysregulation, oxidative phosphorylation, EMT-related pathways such as cadherin switching, and extensive ECM degradation mediated by matrix metalloproteinases (MMPs), collectively reflecting an aggressively proliferative and invasive phenotype. These findings underscore the spatial complexity of tumor ecosystems and highlight the utility of spatially resolved transcriptomics in capturing functionally distinct microenvironments within heterogeneous tumors [32].

**Fig. 4.**
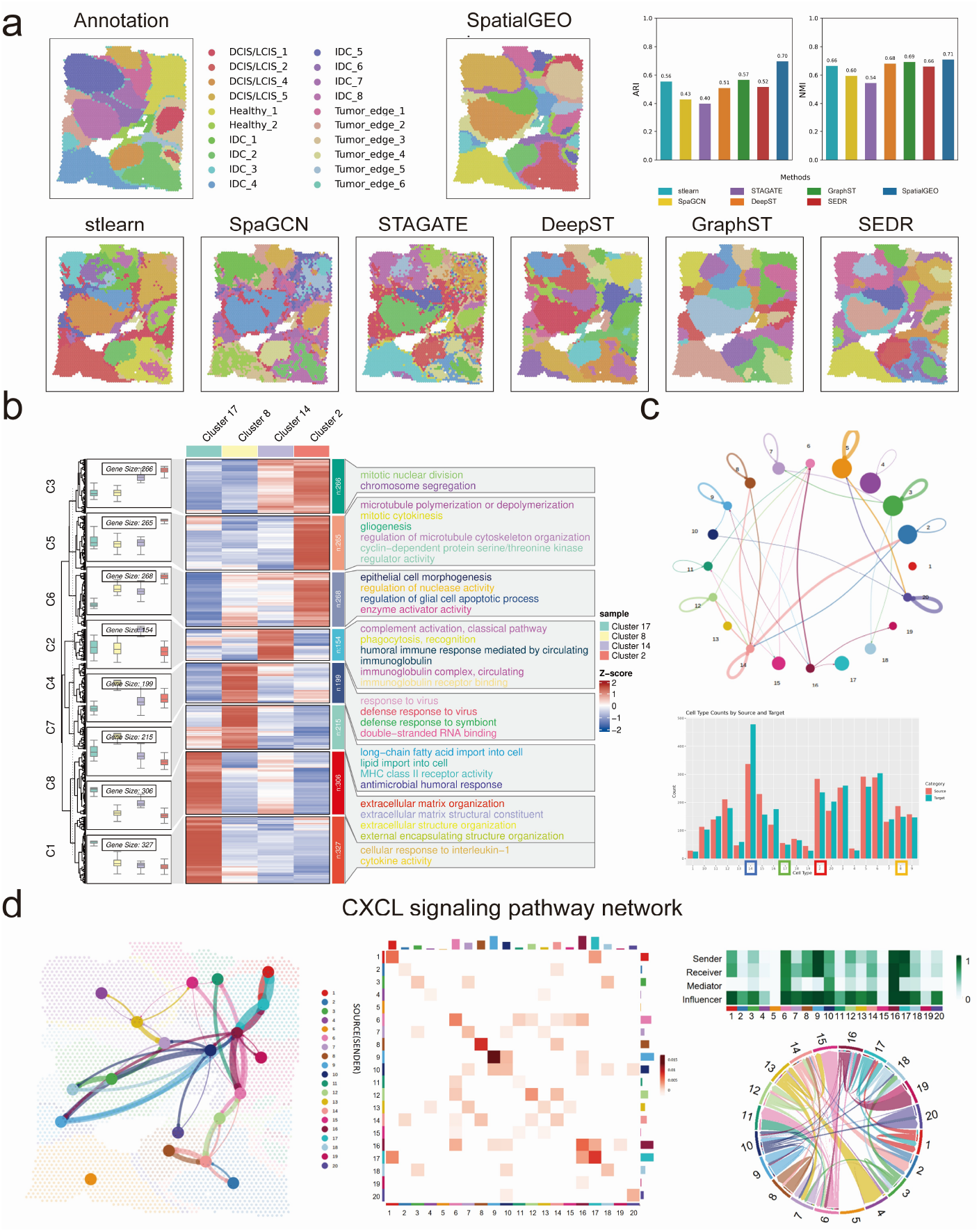
Analysis of breast cancer spatial transcriptomics reveals distinct molecular and cellular features across tumor microenvironments. (**a**) Comparison of clustering performance across methods. Visualization of clustering results from seven methods and bar charts showing the Adjusted Rand Index (ARI) and Normalized Mutual Information (NMI). (**b**) SpatialGEO-identified regions: Cluster17 (healthy tissue), Cluster8 (DCIS/LCIS), Cluster14 (tumor margin), and Cluster2 (IDC). These regions were analyzed using mfuzz clustering and Gene Ontology (GO) enrichment analysis. (**c**) Cell-to-cell communication network, highlighting signaling interactions between distinct tumor microenvironments. (**c**) Results of CXCL pathway communication, illustrating key signaling axes (CXCL8-CXCR1/2 and CXCL13-CXCR5) in breast cancer progression.

**Fig. 5.**
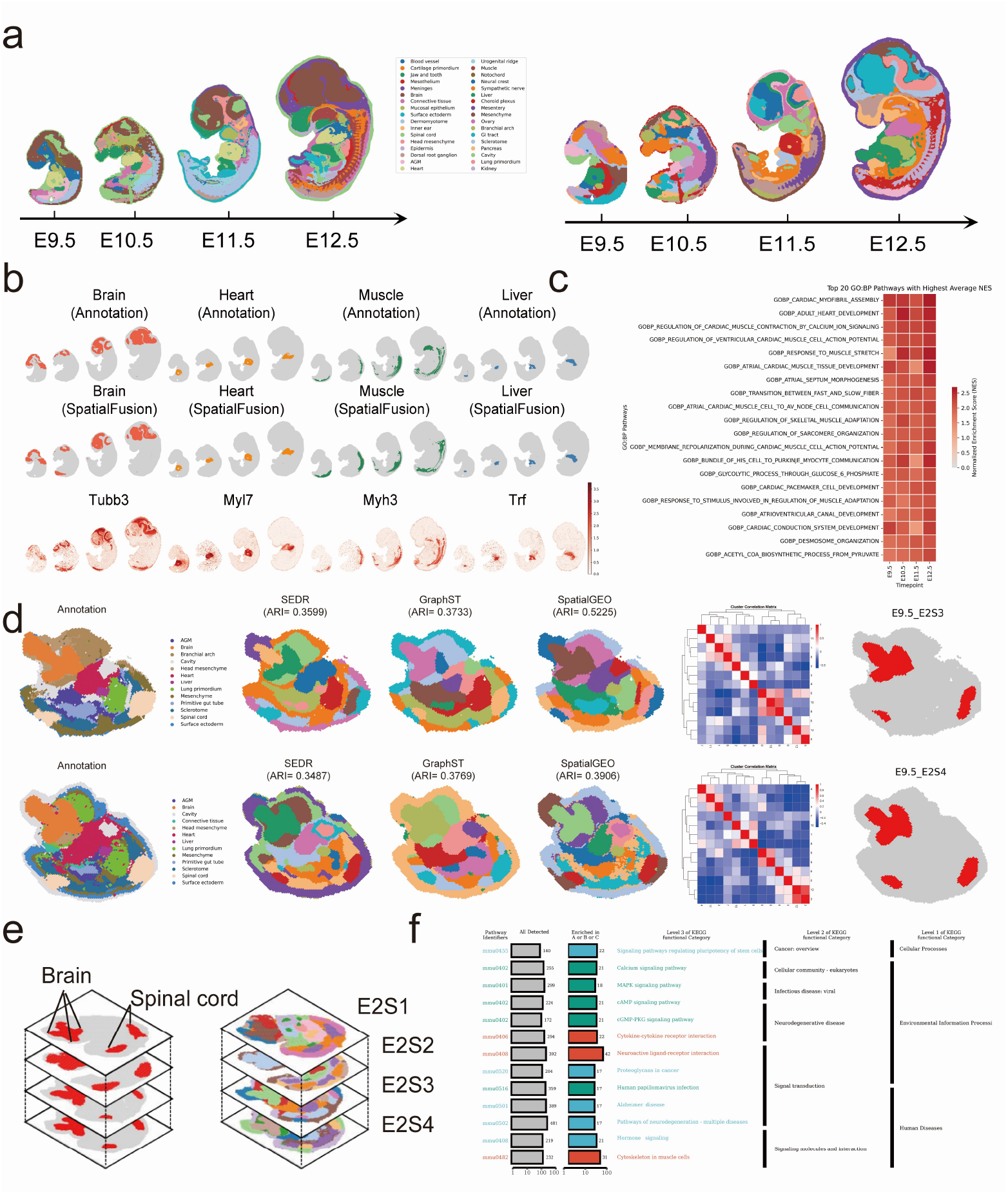
SpatialGEO reveals spatially and temporally resolved transcriptomic domains during mouse embryonic development. (**a**) Annotation of spatial domains across developmental stages in mouse embryonic sections, comparing manual ground truth with domains identified by SpatialGEO. (**b**) Focused analysis of four representative organs—brain, liver, muscle, and heart—across four developmental stages. Rows show, from top to bottom: anatomical annotation, SpatialGEO-predicted domains, expression of canonical marker genes, and module-based expression dynamics, revealing the developmental trajectory of gene expression. (**c**) GSEA enrichment analysis between heart and non-heart regions. The top 20 Gene Ontology biological process (GO BP) terms are shown, ranked by normalized enrichment score (NES). (**d**) (Left) Ground truth annotation of regions E2S3 and E2S4 at stage E9.5, along with spatial domains identified by SEDR, GraphST, and SpatialGEO. (Right) Correlation matrix of SpatialGEO-predicted clusters, highlighting highly correlated regions between brain and spinal cord. (**e**) Spatial domains identified by SpatialGEO across four sections (E2S1–E2S4) (left), and conserved brain–spinal cord relationships across all four sections (right). (**f**) KEGG pathway enrichment analysis of differentially expressed genes between brain and spinal cord regions, combining all four sections.

To further elucidate the functional interactions between spatial domains, we performed cell-to-cell communication analysis using CellChat [33], which revealed distinct signaling behaviors across the identified regions (Figure 5c). In healthy tissue (Cluster17), cells primarily functioned as signaling transmitters, actively secreting chemokines such as CXCL12 [34] and CCL5 to recruit immune effector cells, including T cells and macrophages, thereby maintaining immune surveillance and homeostasis. In contrast, Cluster8 (DCIS/LCIS) exhibited a more balanced signaling profile, functioning as both a signal sender and receiver, and participated in antiviral immune responses and lipid metabolic reprogramming. At the tumor margin (Cluster14), cells predominantly acted as signal receivers, responding to pro-invasive cues—particularly via complement system activation—while also secreting matrix metalloproteinases (MMPs) to promote extracellular matrix (ECM) degradation and facilitate tumor progression. Cluster2 (IDC), by contrast, exhibited a dominant signal-sending profile, relying heavily on autonomous signaling through factors such as vascular endothelial growth factor (VEGF) and MMPs to support unrestrained proliferation, ECM remodeling, and invasion [35].

Among the signaling pathways identified, the CXCL chemokine axis—especially the CXCL8–CXCR1/2 and CXCL13–CXCR5 interactions—emerged as a critical mediator of spatially dependent tumor-immune dynamics (Figure 5d). In Cluster17, the elevated expression of CXCL12, CXCL9 [36], and CXCL10 supported robust immune cell recruitment to the tissue periphery, potentially contributing to early anti-tumor defense. In Cluster8 and Cluster14, bidirectional communication was evident: CXCL12 secreted by Cluster8 may stimulate stromal remodeling at the tumor edge, while Cluster14, in response, activated NF-*κ*B signaling cascades associated with malignant transformation and inflammation. Notably, this reciprocal signaling at the tumor-normal interface reflects a functional switch from immune surveillance to immune evasion and invasion, underscoring the dual role of the CXCL pathway in both tumor suppression and progression.

### 2.5 Spatiotemporal Dissection of the Tissue Microenvironment in Mouse Embryos Using SpatialGEO

To investigate the spatiotemporal heterogeneity of tissue architecture during murine organogenesis, we applied SpatialGEO to four Stereo-seq spatial transcriptomic sections collected at embryonic days E9.5, E10.5, E11.5, and E12.5 (Fig. 5a) [4, 37]. Our analysis focused on four representative organs—brain, heart, liver, and muscle—across these key developmental stages (Fig. 5b). SpatialGEO consistently delineated spatial domains that exhibited high concordance with known anatomical structures, capturing dynamic tissue organization and boundary formation throughout embryogenesis.

To systematically characterize stage-specific regulatory programs during heart development, we conducted gene set enrichment analysis (GSEA) on transcriptional profiles derived from embryonic cardiac tissues at each time point (Fig. 5c). This analysis uncovered a temporally coordinated cascade of biological processes driving cardiac morphogenesis. At E9.5, pathways related to cardiac myofibril assembly and the development of the conduction system were enriched, reflecting the early establishment of cardiac structure and electrical connectivity. By E10.5, gene programs associated with adult heart development, mechanotransduction, and skeletal muscle adaptation were upregulated, indicating the onset of functional maturation and responsiveness to biomechanical cues. At E11.5, enrichment of pathways involved in contraction dynamics and pacemaker cell differentiation suggested the emergence of rhythm-regulating cardiac subpopulations. By E12.5, processes governing Purkinje fiber signaling and membrane repolarization were highly activated, corresponding to the refinement of electrophysiological properties and the consolidation of coordinated heart function. Together, these results delineate a temporally ordered transcriptional program that underlies cardiac morphogenesis, functional specialization, and the maturation of electrophysiological competence during mammalian development.

We also conducted clustering analyses on four spatial transcriptomic sections (E2S1–E2S4) derived from E9.5 mouse embryos to evaluate the performance of SpatialGEO in comparison with SEDR and GraphST (Fig. 5d). SpatialGEO consistently outperformed alternative methods, accurately delineating most anatomically defined regions, as validated against established spatial annotations. Cross-sectional correlation analysis further revealed reproducible expression patterns between anatomically matched brain and spinal cord regions across all sections, reflecting strong biological concordance and developmental symmetry (Fig. 5e). To investigate the molecular programs underlying these spatially coherent domains, we aggregated the corresponding clusters and performed KEGG pathway enrichment analysis on differentially expressed genes (Fig. 5f). This analysis identified significant enrichment in signaling pathways essential for early neural development, including MAPK, calcium, cAMP, and cGMP-PKG signaling. In parallel, pathways associated with cell–cell communication—such as cytokine–cytokine receptor interaction and neuroactive ligand–receptor signaling—were prominently enriched, indicating active intercellular signaling within embryonic neural tissues. Furthermore, pathways regulating cytoskeletal reorganization and muscle contraction were also enriched, consistent with ongoing tissue remodeling and cellular migration during morphogenesis. Remarkably, several pathways commonly implicated in adult neurodegenerative diseases, including Alzheimer’s disease and general neurodegeneration, also emerged as significantly enriched. This suggests that genes traditionally associated with pathological processes may have developmental functions critical for neural circuit formation and maturation. Together, these findings highlight the capacity of SpatialGEO to resolve fine-scale spatial structures and to uncover biologically relevant signaling and structural pathways that govern early neurodevelopmental processes.

## 3 Methods

### 3.1 Spatial transcriptome data processing

#### Data preprocessing

We preprocess Visium spatial transcriptomics data using the *Scanpy* software package. To restrict the analysis to genes with sufficient expression levels, we first filter out lowly expressed genes. Specifically, we retain genes that were expressed in at least 50 cells and have a total count greater than or equal to 10. SpatialGEO then applies a logarithmic transformation to the gene expression matrix to reduce bias and variance, followed by normalization to ensure comparability between spots. For processed data, we employ the *Seurat v3* to select the top *n* highly variable genes (HVGs). The resulting gene expression matrix *X ∈ ℝ*^*m×n*^, where *m* is the number of spatial spots and *n* is the number of selected genes, is used as input to SpatialGEO.

#### Graph construction with spatial transcriptomic data

In order to make full use of the spatial information, SpatialGEO determines the neighbors of a given point according to the Euclidean distance calculated by the spatial position information and selects the nearest *k* points as its neighbors to obtain the adjacency matrix *A ∈* ℝ^*m×m*^. In the adjacency matrix, if the spot *j ∈ V* is a neighbor of the other spot *i ∈ V*, then *a*_*ij*_ = 1; otherwise *a*_*ij*_ = 0. This method of graph construction not only takes into account the physical proximity between spots but also preserves the beneficial properties of spatial information for subsequent tasks. An undirected neighborhood graph *G* = (*V, E*) is constructed based on the adjacency matrix, where *V* denotes the set of spots and *E* denotes the connectivity between points.

### 3.2 Heterogeneous Dual-Source Encoders architecture

#### Autoencoder

The autoencoder is a mapping function consisting of several layers of neural network that maps input data *X ∈* ℝ^*m×n*^ to a latent space representation *Z*_ae_ ∈ ℝ^*m×d*^, where *d ≪ n* is the dimension of the latent space. An encoder contains a number of fully connected layers, each followed by an activation function ReLU, which is used to introduce a nonlinear transformation. The implementation details of the autoencoder are as follows:

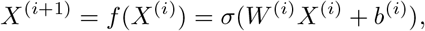

where *W* ^(*i*)^ and *b*^(*i*)^ are the weight and bias of layer *i*, respectively, and *σ* is the LeakyReLU activation function.

The decoder is responsible for mapping the latent representation back to the original feature space and reconstructing the input feature matrix using the following formula:

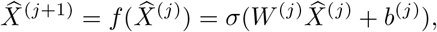

where 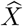 is the reconstructed feature matrix, and *W* ^(*j*)^ and *b*^(*j*)^ are the weights and bias of layer *j*, respectively.

We want to minimise the reconstruction error, that is, to make the reconstructed feature matrix 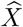 as close as possible to the original input feature matrix *X*. The design loss function is as follows:

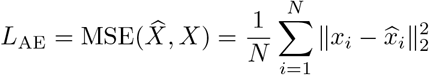

#### Graph Autoencoder

Because the traditional autoencoder is generally symmetric, but the graph neural network is asymmetric. In order to facilitate the fusion of features extracted by autoencoder and graph autoencoder, we improve the traditional graph autoencoder and propose a symmetric graph autoencoder innovatively. The symmetric graph autoencoder maps the input feature matrix *X* and the adjacency matrix *A* to an embedding representation in the potential space *Z*_gae_ ∈ ℝ^*m×d*^, capturing node features and graph structure information. The encoder contains three layers of graph neural network based on the tanh activation function to extract hierarchical information step by step. The potential adjacency matrix is computed by the dot product of *Z*_gae_ and its transpose matrix, followed by the sigmoid activation function 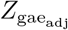, which is used to represent the connectivity relations of the potential graph. A layer of the encoder is expressed as:

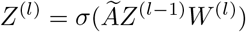

where *W* ^(*l*)^ denotes the learnable parameters of the *l*-th encoder layer.

The decoder reconstructs the input feature matrix and adjacency matrix from the potential space embedding. The decoder also contains three layers of GNN that combine the decoded gene expression matrix 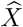 with the adjacency matrix 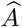. One layer in the decoder is represented as:

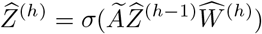

where 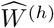 denotes the learnable parameters for the *h*-th decoder layer.

In order to minimise the reconstruction loss function on the weighted attribute matrix and the adjacency matrix, our GAE is designed to minimise the hybrid loss function:

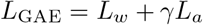

where *γ* is a predefined hyperparameter used to balance the weights of the two reconstruction loss functions. Specifically, *L*_*w*_ and *L*_*a*_ are defined as follows:

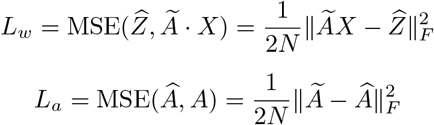

where 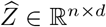 is the reconstructed weighted attribute matrix and 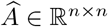 is the reconstructed neighbourhood matrix generated by inner product operation through multilevel representation of the network. By minimizing these equations, the proposed GAE can minimise the reconstruction loss on both the weighted attribute matrix and the adjacency matrix.

### 3.3 Geometry-aware Latent Embedding Representation

First, we combine the potential embeddings of AE (*Z*_ae_ ∈ ℝ^*m×d*^) and GAE (*Z*_sgae_ ∈ ℝ^*m×d*^) with linear combination operations:

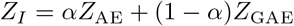

where *d* is the potential embedding dimension and *α* is the learnable coefficient, which is able to selectively determine the importance of two information sources. In SpatialGEO, *α* is initialised to 0.5 and then automatically adjusted using the gradient descent method.

In the process of feature updating, the features of nodes are weighted by the adjacency matrix to capture local neighborhood information. With this operation, we enhance the initial fusion embedding *Z*_*I*_ ∈ ℝ^*m×d*^ by taking into account the local structure within the data:

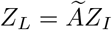

where *Z*_*L*_ *∈ ℝ*^*m×d*^ denotes the local structural enhancement of *Z*_*I*_.

### Geometric Graph Learning

The geometric graph learning architecture consists of multiple graph encoder layers, each of which sequentially performs node aggregation, edge feature updating, and global context conditioning. In this process, the embedding of nodes and edges is gradually refined, eventually generating node representations suitable for a particular task. To ensure stable convergence during the training process, we employ Xavier uniform initialization to initialize the parameters for geometric graph learning. The model can learn node representations that are both expressive and context-aware through geometric graph learning, making full use of global spatial information to achieve robust feature learning.

We use the previously learnt *Z*_*L*_ *∈ ℝ*^*m×d*^ as node features *H*_*v*_ and use the distance between spots as edge features *H*_*E*_. We input the obtained node and edge features into multiple GNN layers for geometric graph learning. To learn multi-scale residual interactions, the node update, edge update, and global context attention modules are employed at the node, edge, and global context levels, respectively.

#### (i) Node Updates

We employ a multi-head attention mechanism for efficient message passing. The feature vectors of node *i* and edge *j → i* in layer *l* are denoted as 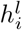 and 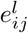, which are converted to a *d*-dimensional space prior to the GNN operation. To update node *i* in layer *l*, we perform message passing as follows:

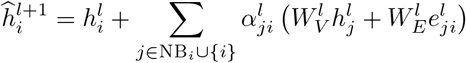

Attention weights 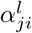 are calculated as follows:

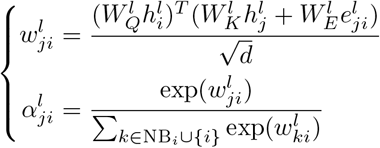

where 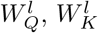, and 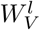 are three weight matrices used to convert the node vectors into query, key, and value representations, respectively. The key and value representations are further complemented by edge vectors using the weight matrix 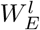. NB_*i*_ represents the neighbors of node *i*. Queries, keys, and values are transformed multiple times to perform parallel attention functions before being concatenated together.

#### (ii) Edge Updates

The edge features *H*_*E*_ are updated through a dedicated MLP-based layer. In this process, the edge features are enriched by concatenating the features of the source and target nodes as well as the original edge features to enrich the edge representation. The concatenated information is processed through a fully connected layer, and after batch normalization and Dropout regularization, the updated edge features are obtained. This mechanism ensures that the edge features can be dynamically updated according to the changes in node features and the connection structure of the graph.

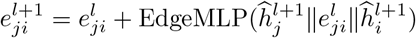

where EdgeMLP denotes the MLP operation for edge updates and ‖ denotes the concatenation operation.

#### (iii) Contextual Feature Extraction

A global context factor *c*^*l*^ is generated by global aggregation of node features *H*_*v*_. The global context factor *c*^*l*^ is used to generate gating signals via an MLP *σ*(GateMLP(*c*^*l*^)) and applied element-wise to the node features, thus regulating the node features to incorporate global information. This process ensures that each node is not only affected by its local neighborhood but is also able to exploit the structure of the entire graph for feature updates, thus generating a more comprehensive node representation.

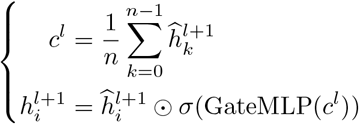

where *n* represents the number of nodes, *σ* denotes the sigmoid function, and GateMLP denotes the MLP for gated attention.

#### (iv) Data Enhancement

To improve the robustness of the model, we introduce a data augmentation strategy during training. Specifically, a small Gaussian noise *ϵ* is added to the node features *H*_*v*_ during training. This enhancement strategy can help the model learn more generalized features and reduce overfitting to specific training instances. In the case of small perturbations in the input data, the model is better able to learn invariant features.

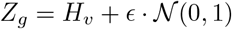

Finally, we obtain the global feature *Z*_*g*_ ∈ *ℝ* ^*m×d*^ that can represent the structure of the global graph.

#### Fusion of Global and Local Features

Finally, we facilitate the efficient transfer of information in the fusion mechanism by introducing jump connections:

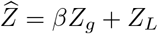

where *β* is a learnable scale parameter. We initialize it to 0 and learn its weights during network training. From a technical point of view, our proposed cross-modal dynamic fusion mechanism combines local and global sample correlations. This design not only finely integrates the information from AE and GAE but also further refines their intrinsic properties to learn potential representations with higher consistency.

#### Target Generation Method

In order to generate more reliable guidance for clustering network training, we first use the more robust clustering embedding 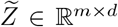, which integrates information from AE and GAE, to generate the target distribution. The generation process consists of two steps:

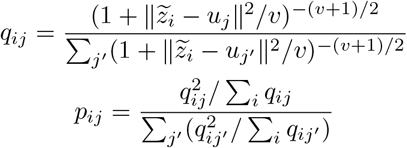

In the first step, we use the Student’s t-distribution to compute the similarity kernel between the *i*-th sample in the fused embedding space 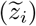 and the *j*-th pre-computed clustering center (*u*_*j*_). *v* is the degree of freedom of the Student t-distribution, and *q*_*ij*_ denotes the probability of assigning the *i*-th node to the *j*-th center. The soft assignment matrix *Q* ∈ *ℝ* ^*m×K*^ reflects the distribution of all samples, and *K* denotes the number of clusters. In the second step, to increase the confidence of the cluster assignment, we introduce a mechanism to drive all samples closer to the cluster centers. Specifically, 0 ≤ *p*_*ij*_ *≤* 1 is an element of the generated target distribution *P* ∈ *ℝ* ^*m×K*^, indicating the probability that the *i*-th sample belongs to the *j*-th clustering center.

Using the iteratively generated target distributions, we compute the soft assignment distributions of AE and GAE on the potential embeddings of the two subnetworks using equation (10), respectively. We denote the soft assignment distributions of GAE and AE as *Q*^*′*^ and *Q*^*′′*^.

In order to train the network in a unified framework and improve the representativeness of each component, we designed the ternary clustering loss by employing a KL divergence of the following form:

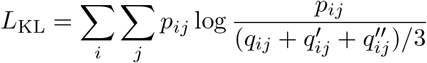

In this formulation, the sum of the soft assignment distributions denoted by AE, GAE, and fusion is simultaneously aligned with the robust target distribution. Since the target distribution is generated without human guidance, we name the loss function ternary clustering loss and the corresponding training mechanism as ternary self-supervised strategy.

#### Joint Loss

The overall learning objective consists of two main components, namely the reconstruction loss for AE and GAE, and the clustering loss associated with the target distribution, which work together to optimize the representation learning and clustering performance of the model. Specifically, the loss function takes the following form:

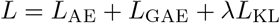

where *λ* is a predefined hyperparameter used to balance the importance between reconstruction loss and clustering loss.

## 4 Discussion

Recent advances in spatial transcriptomics have enabled high-throughput profiling of gene expression while preserving tissue architecture, offering new opportunities to investigate the spatial heterogeneity of complex biological systems.

Here, we propose SpatialGEO, a deep learning framework for spatial geometry awareness, which aims to represent spatial transcriptomic data through feature extraction using a dual-encoder architecture and integrating geometric graph learning. Across datasets spanning diverse biological contexts and spatial resolutions, SpatialGEO consistently outperforms existing methods in spatial domain identification, demonstrating high accuracy and strong concordance with anatomical annotations.

Applying SpatialGEO to human tumour samples reveals its capacity to resolve immune-tumour interfaces with high precision, enabling the identification of localized niches where immune cells interact with malignant populations. This method allows for the delineation of potential immune evasion pathways and uncovers the regulatory dynamics governing immune suppression and activation within the tumour microenvironment. Beyond oncology, SpatialGEO provides a powerful framework for dissecting tissue organization during mammalian development. Analysis of murine embryonic sections demonstrates its ability to distinguish spatial domains associated with distinct developmental stages and tissue types. These spatially resolved profiles capture dynamic changes in gene expression and regulatory programs underlying organogenesis, offering new insights into the spatial and temporal orchestration of developmental processes.

Together, these findings position SpatialGEO as a versatile and scalable tool for interrogating spatial transcriptomic data, with broad applicability to both developmental biology and disease pathology.

## 5 Data availability

The human dorsolateral prefrontal cortex (DLPFC) dataset generated by the Lieber Institute for Brain Development using 10x Visium is available at https://github.com/LieberInstitute/spatialDLPFC. Mouse anterior and posterior brain datasets (10x Visium), as well as the human breast cancer dataset (10x Visium), can be accessed at https://www.10xgenomics.com/resources/datasets. The osmFISH mouse somatosensory cortex dataset is available at http://linnarssonlab.org/osmFISH/. The STARmap mouse visual cortex dataset can be found at https://kangaroo-goby.squarespace.com/data. The Stereo-seq mouse embryo dataset is available via the MOSTA database at https://db.cngb.org/stomics/mosta/.

## 6 Acknowledgements

This work is supported in part by the National Natural Science Foundation of China [62433016, 62202383], and the National Key Research and Development Program of China [2022YFD1801200].

## 7 Author contributions

L.X.Y. designed the overall framework of the SpatialGEO model. J.X.T. developed and implemented the SpatialGEO algorithm. L.X.Y. and J.X.T. enhanced the model design. J.X.T. conducted the majority of the data analyses and generated all figures. Z.D.M. and X.J.L. performed additional data analyses. J.X.T. drafted the manuscript. L.X.Y. and Z.J.N revised and provided feedback. D.G.Y., X.J.L., Q.Y., C.Y.Q., and W.Y.F maintained the computational infrastructure and managed system resources. W.F., S.X.Q. and L.X.Y. conceptualized the project and supervised the overall study. All authors reviewed and approved the final version of the manuscript.

